# Variant calling for cpn60 barcode sequence-based microbiome profiling

**DOI:** 10.1101/749267

**Authors:** Sarah J. Vancuren, Scott J. Dos Santos, Janet E. Hill, the Maternal Microbiome Legacy Project Team

## Abstract

Amplification and sequencing of conserved genetic barcodes such as the cpn60 gene is a common approach to determining the taxonomic composition of microbiomes. Exact sequence variant calling has been proposed as an alternative to previously established methods for aggregation of sequence reads into operational taxonomic units (OTU). We investigated the utility of variant calling for cpn60 barcode sequences and determined the minimum sequence length required to provide species-level resolution. Sequence data from the 5’ region of the cpn60 barcode amplified from the human vaginal microbiome (n=45), and a mock community were used to compare variant calling to de novo assembly of reads, and mapping to a reference sequence database in terms of number of OTU formed, and overall community composition. Variant calling resulted in microbiome profiles that were consistent in apparent composition to those generated with the other methods but with significant logistical advantages. Variant calling is rapid, achieves high resolution of taxa, and does not require reference sequence data. Our results further demonstrate that 150 bp from the 5’ end of the cpn60 barcode sequence is sufficient to provide species-level resolution of microbiota.

## INTRODUCTION

Microbiome profiling is the process of determining which organisms are present in an environment, and their relative abundances. Profiling can be achieved by a “metabarcoding” approach involving targeted PCR and sequencing a genetic barcode: a conserved gene that is shared by many species and can be used to distinguish one from another [1]. Barcodes that have been demonstrated to meet the criteria established by the International Barcode of Life project [2] include the cpn60 gene in bacteria [3], ITS in fungi [4] and cytochrome c oxidase subunit 1 (COI) in animals [2]. Barcodes are important tools for distinguishing species when phenotypic differences are not conclusive; this is especially useful for prokaryotes. Barcode sequences must be universally conserved, so that a wide range of species can be distinguished by the sequence. The cpn60 gene encodes the 60 kDa chaperonin protein conserved in bacteria, eukaryotes and some archaea. A 549-567 bp region of this gene (the “universal target”, UT) has been shown to provide greater resolving power for bacterial species than the widely used 16S rRNA gene sequences [3]. The cpn60 UT sequence can be amplified with universal PCR primers and the chaperonin database, cpnDB, provides a curated collection of chaperonin sequences that can be used for species identification [5,6]. Sequencing of cpn60 UT amplicons from metagenomic samples has been used extensively in microbiome profiling [7–20].

Metagenomic barcode amplicon sequence analysis relies on classifying sequence reads, usually involving formation of operational taxonomic units (OTU), where sequence reads with similarity above a defined threshold are collapsed to a single unit, or other methods to ‘bin’ or categorize reads. De novo assembly of OTU has been particularly useful for cpn60 barcode sequences, allowing a high degree of taxonomic resolution of microbiota [12,13,21,22]. Depending on the assembly method, mapping the raw reads on to the assembled OTU sequences can be subsequently performed to determine OTU abundances [23]. More recently, reference mapping of cpn60 barcode sequence reads was established to speed up processing large amounts of data from microbial communities for which a comprehensive de novo OTU assembly is available for use as a reference, as well as to allow comparison of OTU across studies given a common reference set of OTU. In this procedure, sequence reads are mapped on to the reference sequence database in order to ‘bin’ reads into OTU and calculate their abundances [16,17,24,25].

Formation of exact sequence variants (‘variant calling’) using a denoising algorithm, such as DADA2 for Illumina sequence data, presents another alternative for formation of OTU, by distinguishing sequence errors from true variant positions [26,27]. Potential advantages to this approach include preservation of minor, but informative sequence differences among closely related OTU. Given the resolving power of the cpn60 barcode sequence, variant calling may offer OTU resolution similar to that achieved with de novo assembly, but more rapidly and with lower demand on computational resources. Another significant advantage to this approach is that since exact sequence variants are given a unique identifier based on the actual sequence, it is trivial to recognize identical OTU (variants) across multiple studies.

The objectives of the current study were to determine the length of cpn60 barcode sequence needed for species-level identification, and to compare variant calling to previously established methods for cpn60 OTU generation in terms of logistics, and effects on apparent community composition. Results of this investigation will be applied in ongoing and future studies of microbiomes using the cpn60 barcode sequence.

## METHODS

### Mock community and vaginal microbiome sequencing

A synthetic microbial community (the mixed vaginal panel, MVP) was generated by combining 20 cloned cpn60 barcode sequences as previously described [28]. Plasmid DNA was extracted from *E. coli* with the EZ-10 Spin Column Plasmid DNA Miniprep Kit (Bio Basic Inc., Markham ON, Canada). Purified plasmid DNA was quantified and diluted to 1 × 10^9^ plasmid copies/μL with 10 mM Tris pH 8.5. Five μL of each plasmid solution was pooled to form the synthetic community. cpn60 UT PCR was performed in duplicate with a 1:3 mixture of Illumina adapted primers M729/M280:M1612/M1613 (Table S1) as previously described [29]. Sequence library preparation was conducted according to the Illumina MiSeq 16S Metagenomic Sequencing Library Preparation (Illumina Inc., San Diego, CA) protocol with some minor changes as optimized for the cpn60 UT: in PCR clean-up steps, 32.5 μL of magnetic beads were utilized. Libraries were sequenced on the Illumina MiSeq plaform using a 500 cycle kit with 400 cycles for Read 1 (5’ end of the cpn60 UT) and 100 cycles for Read 2. Only Read 1 sequences were used in downstream analysis. This asymmetric, directional sequencing approach simplifies subsequent bioinformatics steps since the length of the UT sequence prevents complete coverage with a paired end 2 × 250 strategy.

cpn60 UT amplicon sequence data from vaginal microbiome samples (n = 45) generated using the method described above were selected for analysis. Samples were from pregnant women and were selected based on the availability of 75,000 – 96,000 raw reads per sample, providing even levels of input in the different analyses. The original collection of the human microbiome samples was approved by the University of British Columbia Children’s & Women’s Research Ethics Board (certificate no. H17-02253). All sequence data used in comparison of methods is deposited in the National Center for Biotechnology Information (NCBI) Sequence Read Archive and is associated with BioProject accession PRJNA625735.

Cutadapt version 1.15 [30] and trimmomatic version 0.32 (quality score 30, sliding window of 4) [31] were used for amplification primer sequence removal and quality/length trimming, respectively.

### Variant calling

Filtered sequence reads were imported to QIIME2 [32], and DADA2 [27] with a specified truncation length was executed. OTU (variant) sequences were exported from QIIME2 and aligned using watered-BLAST [33] to cpnDB_nr, a non-redundant version of the chaperonin database cpnDB [5,6] to identify the nearest neighbour (NN) for each read. cpnDB_nr consists of cpn60 UT sequences with a single record for each species, and a preference for the type strain if available (downloaded from http://cpndb.ca). Formed OTU sequences that were over 55% identical to anything in cpnDB_nr were considered to be cpn60 sequences and retained for further analysis [34]. The feature table was also exported, and sequence read counts were summed for OTU with the same NN species.

To evaluate an optimal truncation length for the cpn60 UT in terms of the quality of the sequence classification and data retention, truncation lengths of 50, 100, 150, 200, 250 and 300 were tested with DADA2. The OTU sequences formed at each truncation length were aligned to cpnDB_nr with watered-BLAST returning all significant hits [33]. An OTU sequence was considered unambiguously identified if there was only one resulting match to the database that did not tie with any other results.

### De novo assembly

Trinity version 2.4.0 [35] with kmer size of 31 was used to assemble quality filtered reads into OTU. OTU sequence frequencies were determined by mapping quality filtered reads on to the assembled OTU sequences using Bowtie2 version 2.3.3.1 [36]. A feature table including the OTU frequency (read count) in each sample was created by processing the SAM file output from Bowtie2 using components of mPUMA [23]. Assembled OTU sequences were aligned to cpnDB_nr to determine nearest neighbours (NN, best sequence match). For community membership analyses, read counts for individual OTU with the same nearest neighbour species were combined.

### Reference mapping

Filtered reads were mapped to cpnDB_nr or the VOGUE Reference Assembly with Bowtie2 version 2.3.3.1 [36]. The VOGUE Reference Assembly is a curated set of OTU previously generated from a de novo assembly of sequence reads from 546 vaginal microbiomes [14] and labeled according to nearest neighbour in cpnDB_nr. The mock community (MVP) was also mapped on to the 20 cpn60 UT sequences comprising the MVP. Following Bowtie2 mapping to the selected reference database, a feature table containing OTU names and read abundances was created as previously described [17].

### Microbial community composition analysis

Alpha diversity metrics Shannon diversity and Chao1 estimated species richness were calculated for each sample as mean values over 100 subsamples with a sequence depth of 25000, using QIIME2 [32]. Alpha rarefaction plots were visualized to confirm that adequate sampling depth was achieved. The Kruskal-Wallis test by ranks was used to compare diversity and richness metrics resulting from the three methods, and if significant differences were found, *post hoc* pairwise comparisons were done using the Wilcoxon rank sum test.

For initial comparison and visualization of the relationships among microbiome profiles resulting from each OTU generation method in terms of species presence and abundance, a Jensen-Shannon distance matrix was calculated in R using a custom distance function that calculates the square root of the Jensen-Shannon divergence [37]. Hierarchical clustering was conducted using the hclust function in R with the Ward 2 linkage method. Vaginal Community State Type (CST) classification for vaginal samples was based on previously published definitions [14,38,39].

Relationships among microbiome profiles generated for the vaginal samples were also examined by PERMANOVA analysis. Feature tables containing read counts were first subject to centre log-ratio (CLR) transformation to estimate the likelihood of true zero values in the dataset (recommended for sparse, compositional datasets like microbiome profiles) using the aldex.clr function of ALDEx2 [40]. Only OTU present in at least 10% of samples were retained for downstream analysis. CLR-transformed data were converted to a propr object using the aldex2propr function of ALDEx2 and principle component analysis (PCA) was performed on the resulting propr object. PCA data were plotted as compositional biplots (ggbiplot) to visualise similarity of microbiome profiles. To test for significant differences in microbiome profiles by OTU generation method and between samples, the adonis function of vegan R package was used to construct a Euclidean distance matrix from the CLR-transformed data and perform a PERMANOVA on the resultant matrix. As PERMANOVA is sensitive to differences in dispersion of data within groups, it assumes a homogenous within-group dispersion; therefore, we checked this assumption with the betadisper and permutest functions of vegan. The former determines dispersion of data within each group while the latter permutes the dispersion data to test for significant variability of within-group dispersion. Finally, post-hoc testing of permutation data from permutest was performed using the TukeyHSD function to determine where differences in dispersion lay.

## RESULTS AND DISCUSSION

### Optimal length for variant calling

Variant calling using the DADA2 denoising algorithm in QIIME2 employs a truncation length parameter to which all input sequence reads are trimmed uniformly. If a read is under that length then it is not included in the process [27]. It was therefore desirable to determine an optimal truncation length to use with the cpn60 UT sequences, given input reads of up to 400 bp and the trade-off between minimum length requirement, and proportion of reads retained in variant calling.

To investigate the optimal truncation length for cpn60 variant calling with DADA2 without the confounding effect of diminishing read numbers retained as the truncation length requirement increases, we generated a single set of input reads from the vaginal microbiome sequence data that were all at least 300 bp in length after quality filtering. The Trimmomatic quality filtering step was performed so that only reads above 300 bp with a quality threshold of 30 over a 4 bp sliding window would be used as a consistent input for DADA2 (n = 346526 reads). As expected, there were more variants (OTU) produced with the greater sequence truncation lengths (Table 1). This was expected given the way in which DADA2 determines variants; longer lengths mean the algorithm has more chances to detect differences. The OTU identified at each truncation length corresponded to a decreasing number of nearest neighbours as more, lower-quality matches were replaced with fewer, higher quality matches. This observation was also reflected in the increasing proportion of OTU with a single best match. The proportion of OTU unambiguously identified reached a plateau at ∼90% starting at a truncation length of 150 bp (Table 1). Given the protocol with which the samples were sequenced, all reads start from the 5’ end of the cpn60 UT. Any OTU aligned in the reverse direction are therefore considered nonsense and cannot be interpreted for species identification. By 150 bp, 100% of the OTU were aligned in the correct direction (Table 1). Taken together, these results indicate that 150 bp from the 5’ end of the cpn60 barcode sequence is sufficient for microbiome profiling with species-level resolution, consistent with the known features of the cpn60 barcode sequence [3]. Using truncation lengths >150 bp could be desirable depending on the amount of data available, but the obvious trade-off between minimum length and number of reads retained for analysis should be carefully considered. The effect of loss of shorter reads is that the overall sequencing depth of the samples is reduced, and thus the chance of detecting rare organisms, which likely outweighs the benefit of the small increase in resolution that would be achieved using greater lengths. Shorter read lengths could also decrease the cost of sequencing.

**Table 1:** Variant calling results for vaginal microbiome data with different truncations lengths and consistent input of 346526 quality filtered reads.

| Truncation length | Number of OTU | Number of NN | % OTU single hit to cpnDB nr | % NN alignments in the correct direction |
| --- | --- | --- | --- | --- |
| 50 bp | 189 | 150 | 71.4 | 83.6 |
| 100 bp | 186 | 121 | 83.9 | 95.7 |
| 150 bp | 189 | 107 | 90.5 | 100 |
| 200 bp | 197 | 105 | 92.9 | 100 |
| 250 bp | 206 | 105 | 91.3 | 100 |
| 300 bp | 208 | 99 | 93.8 | 100 |

Additional support for the 150 bp length was obtained when we examined the pairwise percent identities calculated from multiple sequence alignments of bacterial sequences from cpnDB_nr (n = 5974), trimmed to lengths ranging from 50 bp to >500 (full length) from the 5’ end (Figure S1). The mean and median percent identities of the pairwise comparisons for alignments of 150 and 200 bp were significantly lower than for longer (or shorter) regions of the cpn60 barcode. This finding is consistent with that of a previous investigation demonstrating relatively higher inter-species sequence differences in the 5’ end of the cpn60 barcode sequence relative to the 3’ end [3].

Considering all the trade-offs of data retention, cost of sequencing, and species resolution, a DADA2 truncation length of 150 bp was used for the subsequent comparison of methods of OTU generation.

### Accuracy of OTU generation methods from synthetic community sequence data

The MVP synthetic community was used to determine accuracy of the three methods of OTU generation. Following amplification primer removal and quality trimming, 54702 and 40128 reads were available for analysis from the duplicate sequencing runs, respectively. The number of OTU produced from de novo assembly, variant calling (truncation length 150), and reference mapping to either the VOGUE reference assembly or MVP sequences was determined (Figure 1).

**Figure 1:**
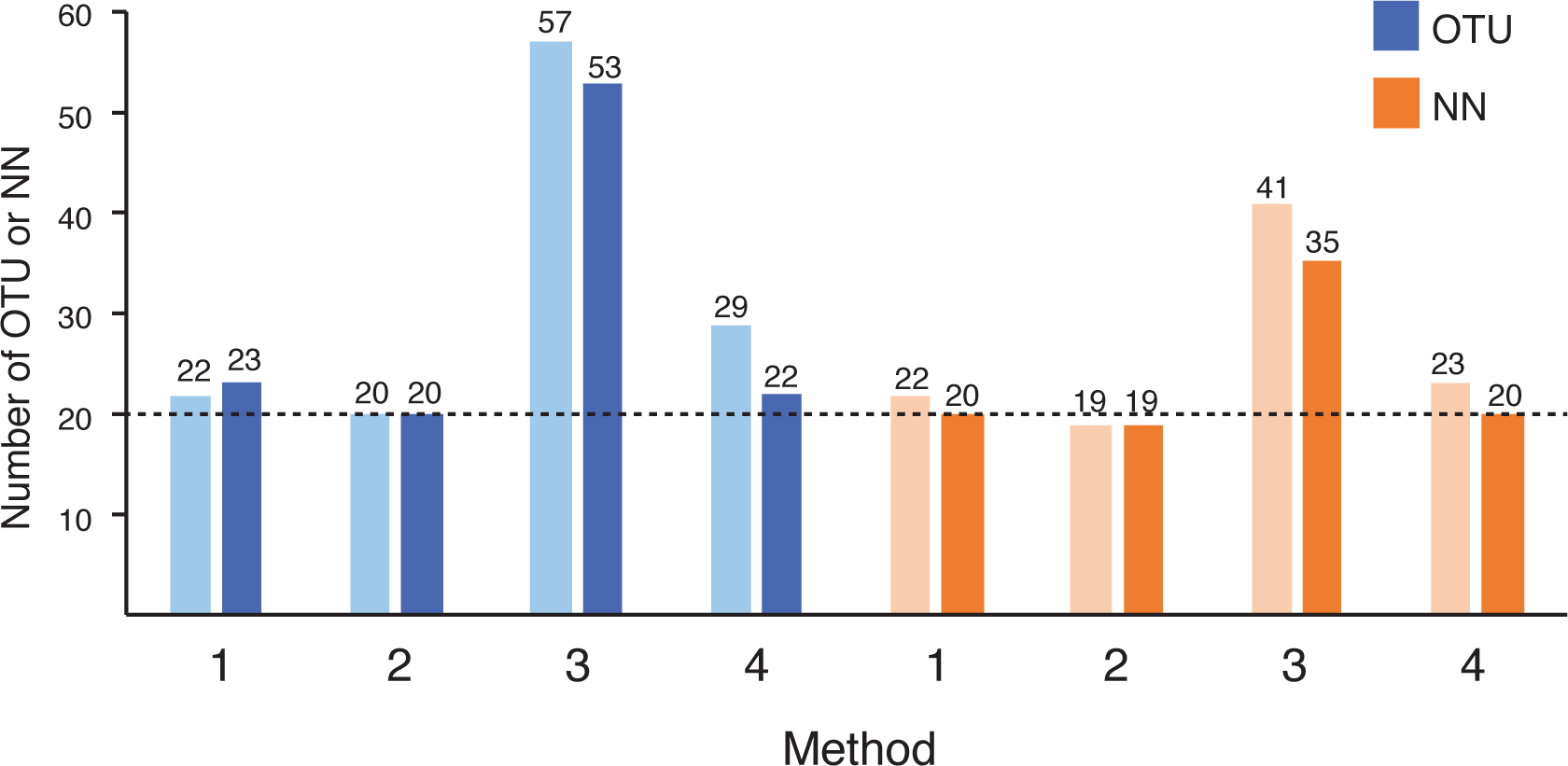
Number of operational taxonomic units (OTU) and nearest neighbour species (NN) from each method applied to the MVP mock community of 20 vaginal bacteria. The synthetic community was sequenced in duplicate in separate runs. For each pair of duplicates, the lighter bar represents the first run and the darker bar the replicate. Methods: 1 = de novo assembly, 2 = reference mapping to MVP, 3 = reference mapping to VOGUE reference assembly, 4 = variant calling.

Reference mapping directly to the MVP gave the expected result of 20 OTU in both library duplicates, confirming that all constituents of the mock community were amplified and sequenced. When the MVP OTU were aligned to cpnDB_nr using watered-BLAST to identify nearest neighbours, they aligned to only 19 different species. This difference was not unexpected since two of the cloned cpn60 barcode sequences in the MVP (corresponding to *Lactobacillus gasseri* and *L. johnsonii*) are 97% identical. Reference mapping to the VOGUE reference assembly was included in the comparison to determine the effect that a larger and more diverse mapping database could have on the results, and we found that far more OTU and NN were generated from the MVP sequence libraries using this approach (Figure 1). Further analysis of the profiles showed that many of the ‘extra’ OTU that were produced were the result of reads mapping to highly similar sequences in the reference database. This result illustrates a limitation of the reference mapping approach, which is dependent on the Bowtie2 algorithm to determine the best match for each read. Bowtie2 alignments close in score may be chosen over one anther unpredictably, or if the program’s search limit has been reached it may not determine a better alignment if one exists, which may lead to changes in resulting profiles [36].

Variant calling and de novo assembly of MVP derived sequence reads both produced more OTU than the 20 expected, up to 29 in the first replicate analysed by variant calling (Figure 1). Additional OTU were most likely formed as a result of minor sequence difference due to PCR errors in the barcode sequence amplification, which was not done with a proof-reading polymerase. Support for this explanation is also found in the observation that the ‘extra’ OTU corresponded to the expected number of nearest neighbours in the second replicate. These observations are typical for descriptions of microbial communities based on barcode amplicon sequencing. OTU formed, especially rare ones, may be either real biological entities or artefacts due to PCR and sequencing errors or the bioinformatic processes used to designate and identify them [41]. We reported all OTU in this analysis regardless of abundance, but a variety of approaches have been applied to curation of OTU, including establishment of cut-offs based on abundance or degree of co-occurrence among samples [42].

Taken together, the results of analysis of the MVP synthetic community sequence data demonstrates that while variant calling and de novo assembly produced the expected community composition, results of reference mapping were heavily influenced by the qualities of the database used and could lead to a significant over-estimation of the number of species in the community.

### Effects of OTU generation method on overall microbial community profiles

Microbial community profiles derived from barcode amplification and sequencing include both an inventory of the distinct OTU present as well as their relative abundance based on sequence read numbers. Various statistical approaches and clustering methods can then be applied to investigate beta-diversity within collections of samples with the goal of identifying similarities and differences among samples that may be associated with defined groups in a study, for example pre- and post-treatment groups, or subjects with different clinical outcomes or phenotypes. To determine the effects of the different OTU formation, identification and frequency calculation methods on overall microbiome profiles, we exploited previously generated data sets derived from the vaginal microbiome (n = 45, 3.8 million raw reads). After quality filtering (q30, minimum length 150), 2 million reads were available for analysis with an average of 44600 reads per sample. For de novo assembly and variant calling, sequence read counts for OTU with identical nearest neighbours in cpnDB_nr were combined to facilitate merging of the resulting read count tables with the output from reference mapping. Results of de novo assembly, reference mapping, and variant calling (using a 150 bp truncation length) were then compared to determine if the methods produce different results in terms of community composition or different conclusions regarding the relationships among samples (alpha and beta diversity).

There was no difference in diversity (Shannon) of the vaginal microbiome profiles determined with the three OTU generation methods (Kruskal-Wallis, *P* > 0.05), but there was a significant difference in richness (Chao1) (Wilcoxon rank sum test, all pairwise comparisons *P* < 0.02) (Figure 2), which was not unexpected given the characteristics of the methods. The relative overestimation of community richness resulting from the reference mapping approach that we observed in the synthetic community sequencing experiment was also observed here. Richness of communities based on OTU generated with variant calling was lower overall than when OTU were generated by de novo assembly, which may reflect the removal of variants determined by the DADA2 denoising algorithm to be the result of sequencing error [26].

**Figure 2:**
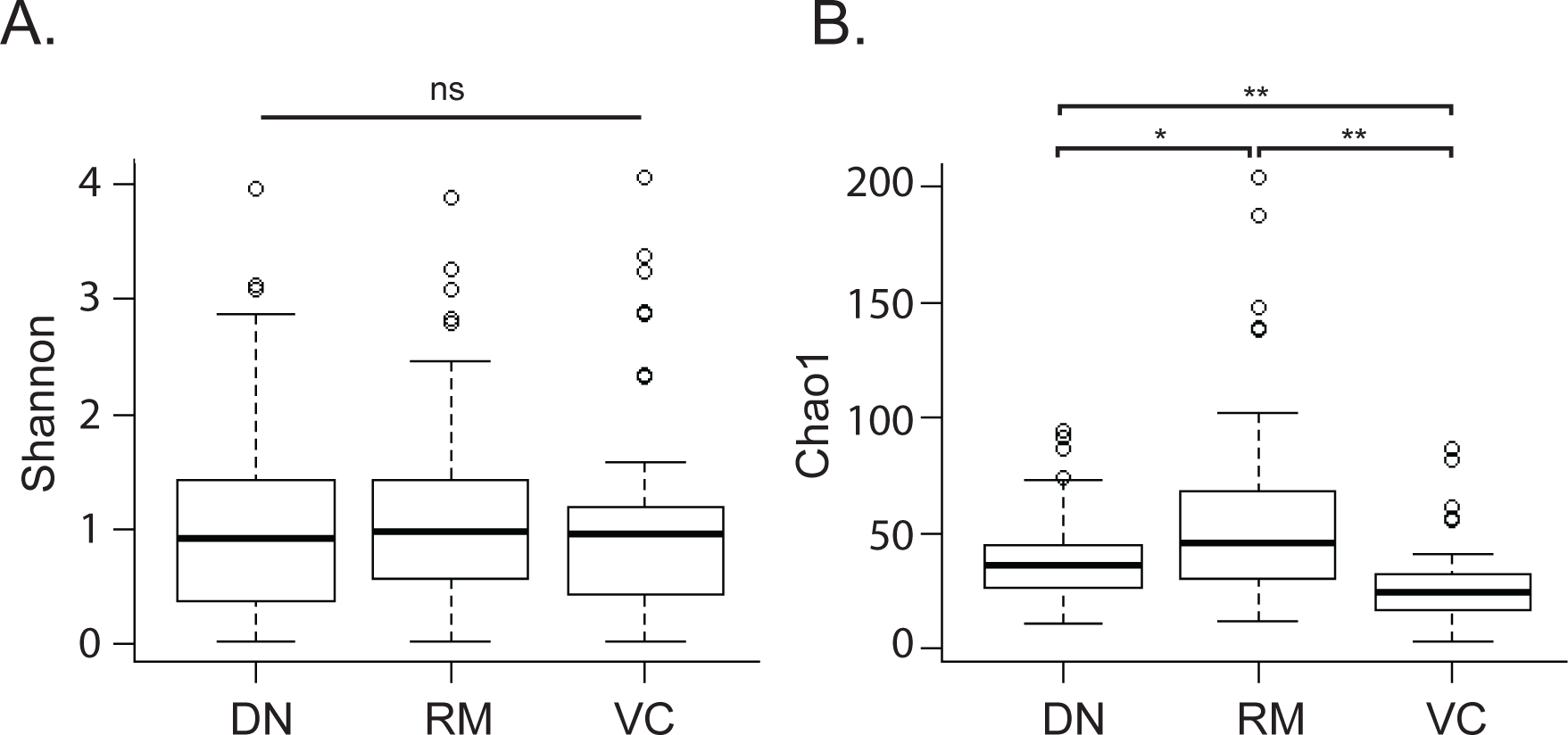
Comparison of alpha diversity metrics. Shannon diversity (A) and Chao1 (B) values were calculated for each vaginal microbiome sample (n = 45) and each method of OTU formation. *P* values are indicated above the bars, * = *P* < 0.02, ** = *P* < 0.001, ns = not significant.

The microbiome profiles of each sample determined with each method were subjected to cluster analysis to determine if the OTU formation method affected the overall microbiome profile and interpretation of the relationships among samples. In most cases, the three profiles for each sample clustered together regardless of OTU generation method (Figure 3). Clusters of samples corresponding to previously defined vaginal microbiome “community state types” (CST) were identified based on visual inspection of proportional abundance of key taxa (Table S2). CST identified were similar to those reported in previous cpn60-based studies of vaginal microbiota of Canadian women: CST I (*Lactobacillus crispatus* dominant), CST II (*Lactobacillus gasseri* dominant or co-dominant), CST III (*Lactobacillus iners* dominant or co-dominant), CST V (*Lactobacillus jensenii* co-dominant), CST IVa (variable, heterogenous mix) and CST IVc (*G. swidsinskii* and/or *G. leopoldii*) [14]. Most (43/45) samples received the same CST classification regardless of the OTU generation method used, demonstrating that one would not make a different conclusion regarding CST affiliation using any of the three methods tested in this study (Figure 3, Table S2). The exceptions were sample numbers 19 and 29. In the results of reference mapping and variant calling for these samples, the most abundant OTU (accounting for 35% and 37% of these samples, respectively) was identified as *Gardnerella leopoldii*, and the second most abundant OTU (29% or 27%) was identified as *Lactobacillus jensenii*, consistent with CST V. Although the proportional abundance of *L. jensenii* was the same in the de novo assembly results for either sample, the most abundant OTU that accounted for 34% and 36% of the microbiome profiles was identified as *G. swidsinskii* (Table S2). The cpn60 sequences of *G. leopoldii* and *G. swidsinskii* are 97% identical to each other. It is surprising that this difference alone would result in a change in the CST identification for these samples and so it is likely that differences in detection of other lower abundance species also contributed.

**Figure 3:**
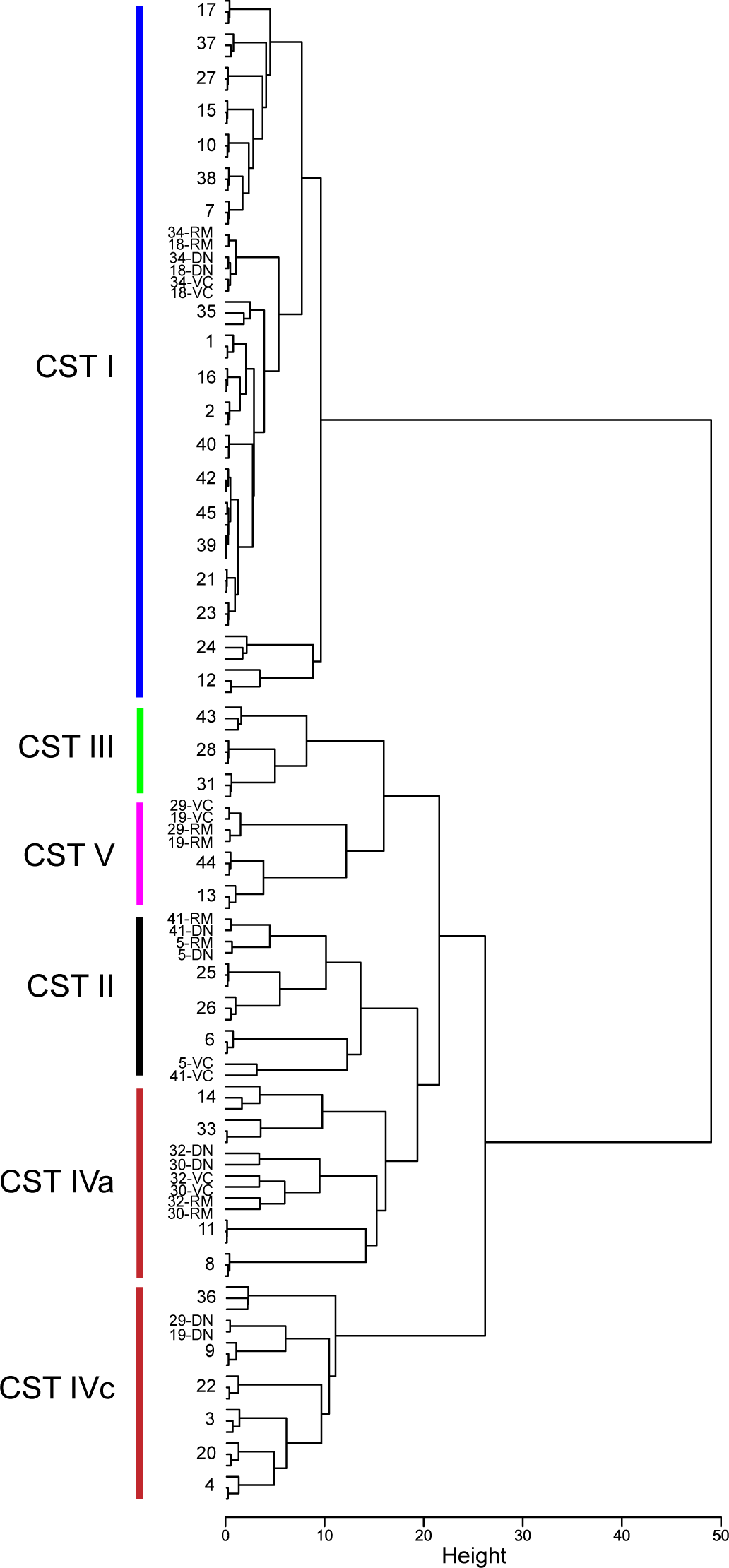
Comparison of microbiome profiles resulting from three methods of OTU generation. Clustering of profiles from vaginal samples based on nearest neighbour species abundance with Jensen-Shannon distance calculation and hclust in R. In cases where the sample profiles from all methods cluster together, only the sample number is shown. RM = reference mapping, DN = de novo assembly, VC = variant calling. Clusters corresponding to CST I-IV are indicated, and are described in the text.

Additional support for the lack of effect of OTU generation method on beta diversity was observed when PERMANOVA analysis was performed. Given that this method is sensitive to group dispersion effects in unbalanced study designs, we also tested for homogeneity of variance. PERMANOVA reported no significant clustering of the vaginal samples by OTU generation method (*P* = 0.986) (Figure 4A); however vaginal microbiome profiles significantly clustered based on sample number (*P* < 0.001) (Figure 4B), meaning that for each sample, the assembly method used to generate the profile did not significantly affect the community composition. No differences in dispersion between groups were seen for either assembly method or sample number (*P* = 0.738 and 0.930, respectively).

**Figure 4:**
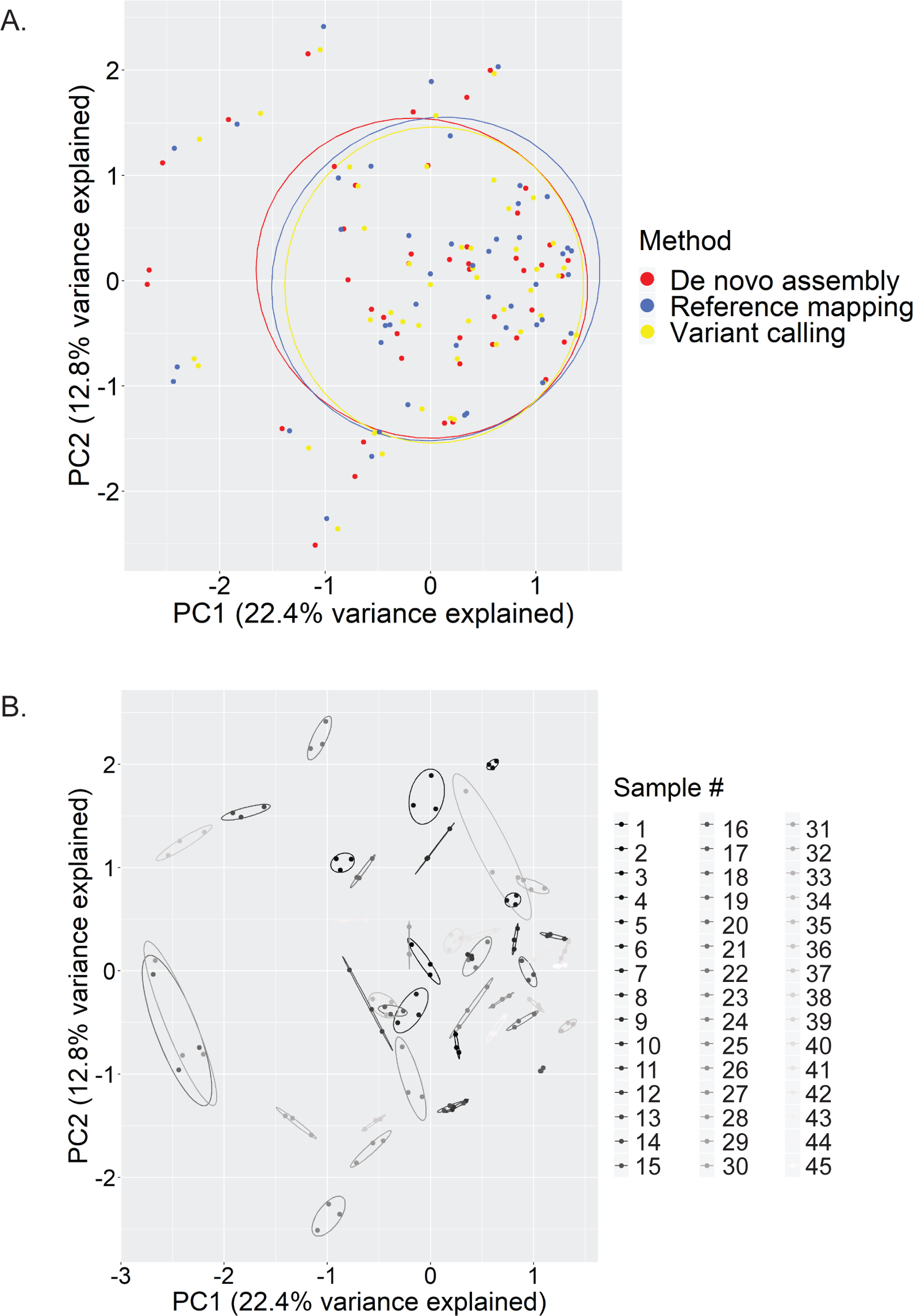
Assembly method does not affect final microbiome composition: Centre log-ratio transformation (CLR) was performed on read count data from all 45 vaginal microbiome profiles generated using three different methods (n = 135). PCA plots grouped by assembly method (A) or sample number (B) were generated and Euclidean distance matrices of CLR-transformed data were analysed by PERMANOVA. Significant clustering of vaginal microbiomes by sample, but not assembly method was evident from PCA plots and supported by PERMANOVA results (*P* < 0.001 and *P* = 0.986, respectively).

### Practical considerations and recommendations

All three approaches to OTU generation use open source software so there is no cost factor to consider. When the real time for all processes in each OTU formation method applied to the 45 vaginal samples (conducted on a Dell PowerEdge R910 with 250 GB RAM running CentOS Linux 7) were summed, reference mapping was the most time-consuming method (58.7 min), followed by de novo assembly (23.7 min) and variant calling (16.1 min). Times for reference-mapping are dramatically increased relative to the other approaches as the size of the reference database increases (data not shown).

The reference mapping approach can be efficient if a reference database customized for the environment under study, but such reference sequence databases are not available for all bacterial communities. The de novo assembly method has been used extensively for cpn60-based microbiome profiling, but is slower, and requires more software dependencies than variant calling. Taken together, the results of the current study suggest that both of these approaches, especially reference mapping, may overestimate microbial community richness. Variant calling was the fastest method we tested, and truncation lengths of at least 150 bp resulted in species level identification of OTU. The use of denoising methods for variant calling has also been suggested for detection of otherwise unobserved 16S rRNA sequence variants [26]. This approach is particularly attractive for users with limited experience with data analysis, since it can be easily installed and accessed within the QIIME2 package [32]. Another significant benefit of variant calling is that unique identifiers are assigned to each variant making it simple to compare detection of variants across studies.

The choice of method for sequence classification from barcode sequence-based studies of microbial communities is influenced by a number of factors including pre-existing knowledge of the environment under study, numbers of samples, sequencing depth, desired level of taxonomic resolution, computational resources, and the need to pool results from independent studies. Exact sequence variant calling with DADA2 using 150 bp from the 5’ end of the cpn60 barcode sequencing is a recommended strategy for rapid, relatively simple to execute, high-resolution profiling of microbiomes that facilitates the comparison of microbiome profiles across studies.

## ACKNOWLEDGEMENTS

The authors are grateful to Champika Fernando for excellent technical assistance.

## AUTHOR CONTRIBUTIONS

Conceived the study: JEH, SJV

Conducted the experiments: SJV, SJD

Analysed the data: SJV, SJD, JEH

Wrote the manuscript: SJV, SJD, JEH

## FUNDING

This work was funded by a Canadian Institutes of Health Research project grant to the Maternal Microbiome Legacy Team, and a Natural Sciences and Engineering Research Council of Canada Discovery Grant to JEH.

## Notes

### Competing Interest Statement

The authors have declared no competing interest.

### Summary of Updates

Demonstration of the recommended variant calling technique and comparison to alternative methods for operational taxonomic unit formation is now presented using a set of 45 human vaginal samples. Figures and supplemental files have been updated.

## REFERENCES

1. Gohl DM, Vangay P, Garbe J, MacLean A, Hauge A, Becker A, et al. Systematic improvement of amplicon marker gene methods for increased accuracy in microbiome studies. Nature Biotechnology. 2016;34: 942–949. doi:10.1038/nbt.3601

2. Hebert PDN, Cywinska A, Ball SL, deWaard JR. Biological identifications through DNA barcodes. Proceedings of the Royal Society B: Biological Sciences. 2003;270: 313–321. doi:10.1098/rspb.2002.2218

3. Links MG, Dumonceaux TJ, Hemmingsen SM, Hill JE. The chaperonin-60 universal target is a barcode for bacteria that enables *de novo* assembly of metagenomic sequence data. PLoS ONE. 2012;7: e49755. doi:10.1371/journal.pone.0049755

4. Schoch CL, Seifert KA, Huhndorf S, Robert V, Spouge JL, Levesque CA, et al. Nuclear ribosomal internal transcribed spacer (ITS) region as a universal DNA barcode marker for Fungi. Proc Natl Acad Sci U S A. 2012;109: 6241–6. doi:10.1073/pnas.1117018109

5. Vancuren SJ, Hill JE. Update on cpnDB: a reference database of chaperonin sequences. Database (Oxford). 2019;2019. doi:10.1093/database/baz033

6. Hill JE, Penny SL, Crowell KG, Goh SH, Hemmingsen SM. cpnDB: a chaperonin sequence database. Genome Research. 2004;14: 1669–75.

7. Hill JE, Seipp RP, Betts M, Hawkins L, Van Kessel AG, Crosby WL, et al. Extensive profiling of a complex microbial community by high-throughput sequencing. Applied and Environmental Microbiology. 2002;68: 3055–66.

8. Dumonceaux TJ, Hill JE, Hemmingsen SM, Van Kessel AG. Characterization of intestinal microbiota and response to dietary virginiamycin supplementation in the broiler chicken. Applied and Environmental Microbiology. 2006;72: 2815–2823.

9. Dumonceaux TJ, Hill JE, Pelletier C, Paice MG, Van Kessel AG, Hemmingsen SM. Molecular characterization of microbial communities in Canadian pulp and paper activated sludge and quantification of a novel *Thiothrix eikelboomii*-like bulking filament. Canadian Journal of Microbiology. 2006;52: 494–500.

10. Desai AR, Musil KM, Carr AP, Hill JE. Characterization and quantification of feline fecal microbiota using cpn60 sequence-based methods and investigation of animal-to-animal variation in microbial population structure. Vet Microbiol. 2009;137: 120–128.

11. Mansfield GS, Desai AR, Nilson SA, Van Kessel AG, Drew MD, Hill JE. Characterization of rainbow trout (*Oncorhynchus mykiss*) intestinal microbiota and inflammatory marker gene expression in a recirculating aquaculture system. Aquaculture. 2010;307: 95–104. doi:Doi 10.1016/J.Aquaculture.2010.07.014

12. Chaban B, Albert A, Links MG, Gardy J, Tang P, Hill JE. Characterization of the upper respiratory tract microbiomes of patients with pandemic H1N1 influenza. PLoS ONE. 2013;8: e69559. doi:10.1371/journal.pone.0069559

13. Chaban B, Links MG, Jayaprakash T, Wagner EC, Bourque DK, Lohn Z, et al. Characterization of the vaginal microbiota of healthy Canadian women through the menstrual cycle. Microbiome. 2014;2: 23. doi:10.1186/2049-2618-2-23

14. Albert AY, Chaban B, Wagner EC, Schellenberg JJ, Links MG, van Schalkwyk J, et al. A study of the vaginal microbiome in healthy Canadian women utilizing cpn60-based molecular profiling reveals distinct *Gardnerella* subgroup community state types. PLoS One. 2015;10: e0135620. doi:10.1371/journal.pone.0135620

15. Bondici VF, Lawrence JR, Khan NH, Hill JE, Yergeau E, Wolfaardt GM, et al. Microbial communities in low permeability, high pH uranium mine tailings: characterization and potential effects. J Appl Microbiol. 2013;114: 1671–1686. doi:10.1111/jam.12180

16. Freitas AC, Chaban B, Bocking A, Rocco M, Yang S, Hill JE, et al. The vaginal microbiome of healthy pregnant women is less rich and diverse with lower prevalence of Mollicutes compared to healthy non-pregnant women. Scientific Reports. 2017;7: 9212. doi:10.1038/s41598-017-07790-9

17. Freitas AC, Bocking A, Hill JE, Money DM, Money D, Bocking A, et al. Increased richness and diversity of the vaginal microbiota and spontaneous preterm birth. Microbiome. 2018;6: 117. doi:10.1186/s40168-018-0502-8

18. Oliver KL, Hamelin RC, Hintz WE. Effects of transgenic hybrid aspen overexpressing polyphenol oxidase on rhizosphere diversity. Appl Environ Microbiol. 2008;74: 5340–8.

19. Peterson SW, Knox NC, Golding GR, Tyler SD, Tyler AD, Mabon P, et al. A Study of the Infant Nasal Microbiome Development over the First Year of Life and in Relation to Their Primary Adult Caregivers Using cpn60 Universal Target (UT) as a Phylogenetic Marker. PLoS ONE. 2016;11: e0152493. doi:10.1371/journal.pone.0152493

20. McKenney EA, Ashwell M, Lambert JE, Fellner V. Fecal microbial diversity and putative function in captive western lowland gorillas (*Gorilla gorilla gorilla*), common chimpanzees (*Pan troglodytes*), Hamadryas baboons (*Papio hamadryas*) and binturongs (*Arctictis binturong*). Integrative Zoology. 2014;9: 557–569. doi:10.1111/1749-4877.12112

21. Costa MO, Chaban B, Harding JCS, Hill JE. Characterization of the fecal microbiota of pigs before and after inoculation with “*Brachyspira hampsonii*.” PLoS ONE. 2014;9: e106399.

22. Links MG, Demeke T, Gräfenhan T, Hill JE, Hemmingsen SM, Dumonceaux TJ. Simultaneous profiling of seed-associated bacteria and fungi reveals antagonistic interactions between microorganisms within a shared epiphytic microbiome on *Triticum* and *Brassica* seeds. New Phytologist. 2014;202: 542–553.

23. Links MG, Chaban B, Hemmingsen SM, Muirhead K, Hill JE. mPUMA: a computational approach to microbiota analysis by *de novo* assembly of OTUs based on protein-coding barcode sequences. Microbiome. 2013;1: 23. doi:10.1186/2049-2618-1-23

24. Links MG, Chaban B, Hemmingsen SM, Muirhead K, Hill JE. mPUMA: a computational approach to microbiota analysis by de novo assembly of operational taxonomic units based on protein-coding barcode sequences. Microbiome. 2013;1: 23. doi:10.1186/2049-2618-1-23

25. Albert AYK, Chaban B, Wagner EC, Schellenberg JJ, Links MG, van Schalkwyk J, et al. A Study of the Vaginal Microbiome in Healthy Canadian Women Utilizing cpn60-Based Molecular Profiling Reveals Distinct Gardnerella Subgroup Community State Types. Fredricks DN, editor. PLOS ONE. 2015;10: e0135620. doi:10.1371/journal.pone.0135620

26. Callahan BJ, McMurdie PJ, Holmes SP. Exact sequence variants should replace operational taxonomic units in marker-gene data analysis. The ISME Journal. 2017;11: 2639–2643. doi:10.1038/ismej.2017.119

27. Callahan BJ, McMurdie PJ, Rosen MJ, Han AW, Johnson AJA, Holmes SP. DADA2: High-resolution sample inference from Illumina amplicon data. Nature Methods. 2016;13: 581–583. doi:10.1038/nmeth.3869

28. Dumonceaux TJ, Schellenberg J, Goleski V, Hill JE, Jaoko W, Kimani J, et al. Multiplex detection of bacteria associated with normal microbiota and with bacterial vaginosis in vaginal swabs using oligonucleotide-coupled fluorescent microspheres. Journal of Clinical Microbiology. 2009;47: 4067–4077. doi:10.1128/jcm.00112-09

29. Hill JE, Town JR, Hemmingsen SM. Improved template representation in *cpn*60 PCR product libraries generated from complex templates by application of a specific mixture of PCR primers. Environmental Microbiology. 2006;8: 741–746. doi:doi: 10.1111

30. Martin M. Cutadapt removes adapter sequences from high-throughput sequencing reads. EMBnet.journal. 2011;17: 10–12. doi:10.14806/ej.17.1.200

31. Bolger AM, Lohse M, Usadel B. Trimmomatic: a flexible trimmer for Illumina sequence data. Bioinformatics. 2014;30: 2114–20. doi:10.1093/bioinformatics/btu170

32. Bolyen E, Rideout JR, Dillon MR, Bokulich NA, Abnet CC, Al-Ghalith GA, et al. Reproducible, interactive, scalable and extensible microbiome data science using QIIME 2. Nat Biotechnol. 2019;37: 852–857. doi:10.1038/s41587-019-0209-9

33. Schellenberg J, Links MG, Hill JE, Dumonceaux TJ, Peters GA, Tyler S, et al. Pyrosequencing of the chaperonin-60 universal target as a tool for determining microbial community composition. Applied and Environmental Microbiology. 2009;75: 2889–2898.

34. Johnson LA, Chaban B, Harding JC, Hill JE. Optimizing a PCR protocol for cpn60-based microbiome profiling of samples variously contaminated with host genomic DNA. BMC Research Notes. 2015;8: 253. doi:10.1186/s13104-015-1170-4

35. Grabherr MG, Haas BJ, Yassour M, Levin JZ, Thompson DA, Amit I, et al. Full-length transcriptome assembly from RNA-Seq data without a reference genome. Nature Biotechnology. 2011;29: 644–52. doi:10.1038/nbt.1883

36. Langmead B, Salzberg SL. Fast gapped-read alignment with Bowtie 2. Nature Methods. 2012;9: 357–9. doi:10.1038/nmeth.1923

37. Endres DM, Schindelin JE. A new metric for probability distributions. IEEE Trans Inf Theory. 2003;49: 1858–1860.

38. Ravel J, Gajer P, Abdo Z, Schneider GM, Koenig SS, McCulle SL, et al. Vaginal microbiome of reproductive-age women. Proceedings of the National Academy of Sciences of the United States of America. 2011;108 Suppl 1: 4680–7. doi:1002611107 [pii] 10.1073/pnas.1002611107

39. Gajer P, Brotman RM, Bai G, Sakamoto J, Schutte UM, Zhong X, et al. Temporal dynamics of the human vaginal microbiota. Science Translational Medicine. 2012;4: 132ra52. doi:10.1126/scitranslmed.3003605

40. Fernandes AD, Macklaim JM, Linn TG, Reid G, Gloor GB. ANOVA-Like Differential Expression (ALDEx) Analysis for Mixed Population RNA-Seq. Parkinson J, editor. PLoS ONE. 2013;8: e67019. doi:10.1371/journal.pone.0067019

41. Kunin V, Engelbrektson A, Ochman H, Hugenholtz P. Wrinkles in the rare biosphere: pyrosequencing errors can lead to artificial inflation of diversity estimates. Environmental Microbiology. 2010;12: 118–123. doi:10.1111/j.1462-2920.2009.02051.x

42. Frøslev TG, Kjøller R, Bruun HH, Ejrnæs R, Brunbjerg AK, Pietroni C, et al. Algorithm for post-clustering curation of DNA amplicon data yields reliable biodiversity estimates. Nat Commun. 2017;8: 1188. doi:10.1038/s41467-017-01312-x

