## Supplementary material for "Variant calling for cpn60 barcode sequence-based microbiome profiling": Figure S1

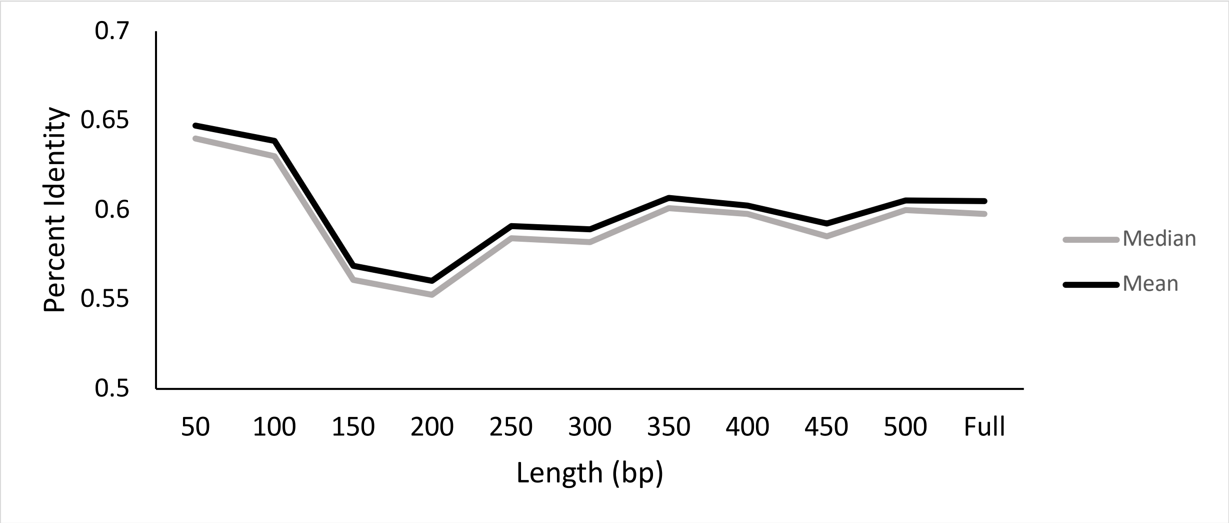


**Figure S1: Multiple All vs. All bacterial cpnDB_nr alignments.** Various lengths starting from the 5' end of the *cpn*60 UT were extracted from a bacteria-only version of cpnDB_nr. Alignments were conducted with clustalw2 and a similarity table was formed with dnadist from the phylip package. Mean and median percent identities were determined for each set of alignments.
