## Supplementary material for "Variant calling for cpn60 barcode sequence-based microbiome profiling": Table S1

**Table S1: Illumina adapted *cpn*60 PCR primer sequences.**

| Primer | Sequence (5’-3’) |
|---|---|
| M279 | TCG TCG GCA GCG TCA GAT GTG TAT AAG AGA CAG GAI III GCI GGI GAY GGI ACI ACI AC |
| M280 | GCT TCG TGG GCT CGG AGA TGT GTA TAA GAG ACA GYK IYK ITC ICC RAA ICC IGG IGC YTT |
| M1612 | TCG TCG GCA GCG TCA GAT GTG TAT AAG AGA CAG GAI III GCI GGY GAC GGY ACS ACS AC |
| M1613 | GCT TCG TGG GCT CGG AGA TGT GTA TAA GAG ACA GCG RCG RTC RCC GAA GCC SGG IGC CTT |
